# The Broken Window: An algorithm for quantifying and characterizing misleading trajectories in ecological processes

**DOI:** 10.1101/2020.07.07.192211

**Authors:** Christie A. Bahlai, Easton R. White, Julia D. Perrone, Sarah Cusser, Kaitlin Stack Whitney

## Abstract

A core issue in temporal ecology is the concept of trajectory—that is, when can ecologists have reasonable assurance that they know where a system is going? In this paper, we describe a *non-random resampling* method to directly address the temporal aspects of scaling ecological observations by leveraging existing data. Findings from long-term research sites have been hugely influential in ecology because of their unprecedented longitudinal perspective, yet short-term studies more consistent with typical grant cycles and graduate programs are still the norm. We use long-term insights to create ‘broken windows,’ that is, reanalyze long-term studies from short-term observational perspectives to examine discontinuities in trends at differing temporal scales.

The broken window algorithm connects our observations between the short-term and the long-term with an automated, systematic resampling approach: in short, we repeatedly ‘sample’ moving windows of data from existing long-term time series, and analyze these sampled data as if they represented the entire dataset. We then compile typical statistics used to describe the relationship in the sampled data, through repeated samplings, and then use these derived data to gain insights to the questions: 1) *how often are the trends observed in short-term data misleading, and* 2) *can characteristics of these trends be used to predict our likelihood of being misled?* We develop a systematic resampling approach, the ‘broken_window algorithm, and illustrate its utility with a case study of firefly observations produced at the Kellogg Biological Station Long-Term Ecological Research Site (KBS LTER). Through a variety of visualizations, summary statistics, and downstream analyses, we provide a standardized approach to evaluating the trajectory of a system, the amount of observation required to find a meaningful trajectory in similar systems, and a means of evaluating our confidence in our conclusions.

**Highlights:** Trends identified in short-term ecology studies can be misleading.

Non-random resampling can show how prone different systems are to misleading trends

The Broken Window algorithm is a new tool to help synthesize temporal data

This tool helps to understand how much data is needed for forecasting to be reliable It can also be used to quantify how likely it is that an observed trend is spurious.

## 1 Introduction

A fundamental problem in ecology is understanding how to scale discoveries: from patterns observed in the lab or the plot to the field or the region, or bridging between short-term observations to long term trends and trajectories (Chave, 2013; Levin, 1992; Schneider, 2001). While shorter-term studies (i.e. those where data collection occurs for less than ∼5 years) that coincide with length of typical grant cycles and graduate programs are still the norm, these human constraints do not necessarily capture the ecological phenomena they seek to measure, particularly their temporal dependencies (Hastings, 2004; Wood et al., 2020). This unfortunate mismatch of scales has the potential to limit our understanding of ecological trajectories-that is, the direction a system is going through time, and can undermine our efforts towards a predictive ecology (Evans et al., 2012). Understanding where and how short term patterns fit into broader trajectories, and how to interpret short-term patterns in the context of a system’s trajectory remains an open question (Wauchope et al., 2019; White, 2019). This is illustrated by the recent insect decline controversy, where several high profile papers have observed precipitous declines in insect populations have been subsequently shown to use inappropriate methods for synthesizing the data (Daskalova et al., 2021; Grames et al., 2019; D. L. Wagner, 2020). For example, it is inappropriate to simply combine multiple short term studies and extrapolate (e.g.: Sánchez-Bayo & Wyckhuys, 2019), particularly without explicitly considering the underlying temporal dependencies in the data (Didham et al., 2020).

However, simply recommending that scientists collect more data, for longer, is not necessarily practicable. While long term studies are hugely influential in ecology, they require long-term access to research resources and infrastructure and thus their unprecedented longitudinal perspective is not typical (Hughes et al., 2017). Furthermore, short-term studies, given their prevalence and more limited temporal commitments, can provide a more spatially distributed and potentially richer and more nuanced view into a specific phenomenon at a point in (or shorter period of) time. The key to meaningful synthesis of this vast resource of short-term studies is linking the two extremes of scale. Thus, long-term data, particularly those produced in networked, uniform approaches like those offered by the United States Long Term Ecological Research Network (LTER), present a fundamental opportunity to bridge short and long-term trends through data mining. With long term data, ecologists can systematically investigate the presence and prevalence of short-term trends and compare them to the long-term system trajectories these data document.

Ecological systems are inherently dynamic, and variations in the metrics humans collect about these systems can be driven by a variety of stochastic and deterministic processes, as well as by sampling error or other research-induced effects (Suding & Gross, 2006). Short-term dynamics observed in an ecological system are not always indicative of the long-term trajectory of that system (Carey & Cottingham, 2016), and furthermore, shorter observation periods can lead to spurious observations because of sampling error variance (Daskalova et al., 2021). In population processes, for example, density-dependent deterministic mechanisms, combined with environmental perturbations, can produce highly variable population numbers over various time scales (Turchin, 2003). Decoupling these processes can reveal the skeleton of a deterministic process interacting with external forces (Bahlai & Zipkin, 2020). However, to disentangle these drivers from an empirical standpoint generally requires a substantial amount of data to be collected over time (Cusser et al., 2020; Higgins et al., 1997). Indeed, in a recent study, White (2019) found that 72% of vertebrate population monitoring programs required at least a decade of observation before the overall trajectory of the population could be detected statistically. A recent study of trends in water bird populations found that short term trends were generally reflective of longer-term patterns (Wauchope et al., 2019), but varied by the generation length of the organism under study. However, they found that, similar to the White (2019) study, greater than two decades of observations would be required to reliably detect a change of 1% per year. Conversely, a study of population viability modelling in snails determined that although longer time series were generally better for establishing the population’s trajectory, diminishing returns in precision were observed after about 10-15 years of data were collected (Rueda-Cediel et al., 2015). It is unclear how these findings can be generalized across organisms with differing lifespans, reproductive strategies and life histories, or other environmental processes.

The question of trajectory over time is central in ecology, particularly as related to how ecological systems on which humans depend are responding to disturbance or will behave under future climate or environmental conditions (Sutherland et al., 2013). Trajectory is essential to our understanding of ecosystems, their management, and policy decisions, as we interact with our environment. Analytic approaches to time series data have long been a focal area of research in ecology, allowing practitioners to examine temporal dependencies in a variety of processes. The shape a time series takes can provide meaningful information about the properties of the system, the rules that govern its variability, and the trajectory that the system is taking (Esling & Agon, 2012). But when insufficient data exists to apply (or even select) an appropriate time-series approach, a scientist may resort to simpler statistical tools, such as linear models, to describe the patterns observed in the data through the study’s window of observation. It is not uncommon for a shorter-duration multi-year ecological study to extrapolate from its data, using the trends observed within their sampling window to draw conclusions about a system’s apparent trajectory. For example, a study of British ladybeetle communities concluded that native ladybeetle species were in decline, as was total ladybeetle abundance, following the introduction of an invasive species (Brown et al., 2011). Another found that the richness and abundance of seeds in a soil seed bank were in a recovery trajectory following a period of industrial pollution (M. Wagner et al., 2006). An adventive pest species was implicated in reducing carbon to nitrogen ratios, organic matter in soils of infested forests, thus substantially changing the ecosystem’s function over time (Orwig et al., 2008). These examples, representing very different ecological domains, have a common element of a three-year study duration. Yet these inferences may be out of temporal sync with the processes they aim to understand (Birkhead, 2014).

A vexing problem arises when shorter term studies apply statistical tools at time scales that are not matched with the underlying processes to make inferences about trajectory: not only may spurious trends be observed, but because only a portion of the underlying process variability is captured, a higher degree of statistical confidence in the result will be found. For example, Bahlai and students examined a 12-year time series of firefly captures from Michigan (Hermann et al., 2016). Concerns had been raised about the status of fireflies in eastern North America (Chow et al., 2014), however, for that population, the authors found no evidence of decline over the 12 years (**Fig. 1**): there was no linear relationship between average captures and year (*p*=0.71, R^2^=0.002), and, indeed, there appeared to be evidence of a cyclical dynamic common to many populations near their carrying capacity (**Fig. 1A**). However, students remarked that if the study had been limited to, for example, the 4 years from 2005-2008 (**Fig. 1B**), dramatically different conclusions would have been made. A linear regression of these data would very likely have been interpreted as ‘strong evidence’ that a decline was occurring in this population (slope - 0.31±0.05, p=0.000003, R^2^=0.633). Simply, with less data, we would have made the wrong conclusions, and we would have been very confident in our wrong answer. This connection between shorter observation periods and more pronounced patterns is supported by observations made in synthesis efforts: in a compilation of insect biodiversity studies, the shortest time series were more likely to show the most extreme trends (Daskalova et al., 2021).

**Figure 1:**
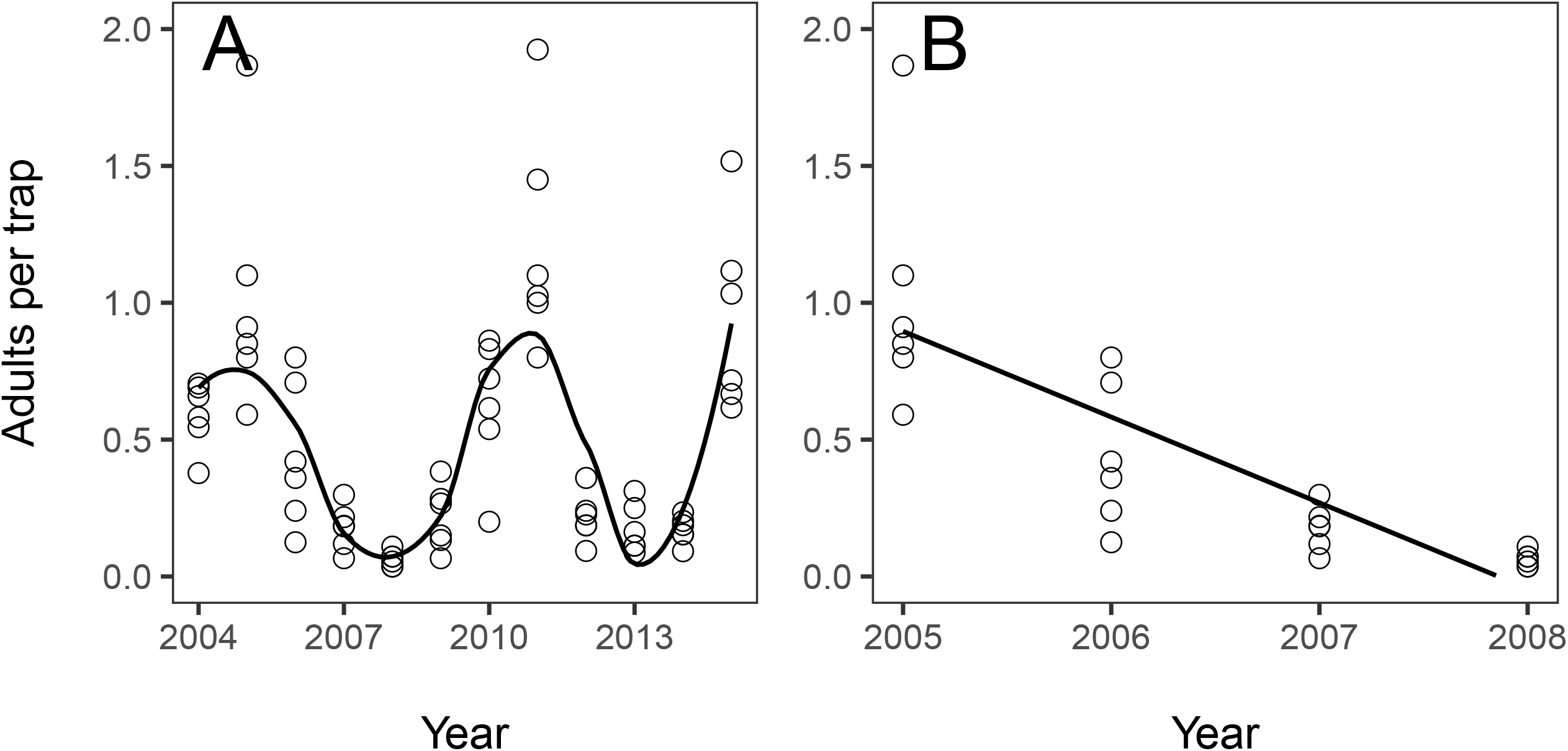
Same data, different observation periods, different conclusions. Firefly populations (reported as mean number of adults captured per trap) monitored in ten plant community treatments at Kellogg Biological Station in southwestern Michigan cycle over an approximately 6 year period (panel A). Yet, if sampling had only occurred over a 4 year period, we would conclude the population underwent a steep (and statistically significant) decline in the four years from 2005-2008 (slope -0.31±0.05, p=0.000003, R^2^=0.633; panel B). Data and figures adapted from Hermann et al (2016).

It is because of this phenomenon of “highly-confident wrong answers” that long-term studies are so valued in the ecological community. Indeed, because biological systems are often defined by their variability, when studies are shown to be irreproducible, it is not necessarily due to poor research practice, but due to their inability to capture the full variability of the system within the limits of the study design (Jarvis & Williams, 2016; Voelkl & Würbel, 2016). Long-term ecological research provides insight into the inherent variability of natural systems (Lovett et al., 2007), and insights are thus often only apparent after many years of study (Knapp et al., 2012). Beyond this, there are many other inherent benefits to long-term studies. Long-term studies are disproportionately represented in policy reports and in the ecological literature: studies involving long term observations are cited more often than studies of shorter duration (Hughes et al., 2017). Furthermore, long-term observational studies provide important baseline data: as the world itself changes, these data provide insight into how ecosystems function, instead of studying phenomena after they happen (Franklin et al., 1990; Hastings, 2004; Magurran et al., 2010).

Although the importance of long-term studies is clear, empirical examinations of the converse are rare: just how frequently are scientists misled by short-term studies? Can knowledge generated by studying the relationship between short- and long-term studies to bridge the interpretations of short-term data to long-term processes? In this study, we describe a synthetic, computational approach to create a framework to address two hypotheses:

*Shorter observation periods will increase the likelihood of observing misleading trends*

> Because exogenous forces are of greater influence at smaller spatial and temporal scales, we predict that short time periods will be more variable due to these processes, and conversely do not capture the full extent of natural variability (Lovett et al., 2007; Suding & Gross, 2006), so they are more likely to result in “highly-confident wrong answers.”

*Statistical metrics often used as a proxy for ‘confidence’ in short-term trends (such as the p-value) will not be associated with an increased likelihood of capturing a time period consistent with long-term trends*.

> Following from the previous prediction, we predict that p-values will be inferior predictors of the ‘correctness’ of short-term trends in predicting longer term trajectory compared to other properties of the system. Better predictors may include statistical measures (slope, standard error), but trends are likely moderated by system specific predictors (e.g. site, data type).

The Broken Window Algorithm is a suite of tools which will allow ecologists to leverage existing data to make inferences about system behavior and data needs to characterize system trajectories using an automated, non-random resampling approach (White & Bahlai, 2020): in short, our algorithm repeatedly ‘samples’ sequential moving windows of data from existing long-term time series, and analyzes these sampled data as if they represented the entire dataset that is, using knowingly limited ‘windows’ of observation to determine how temporal dependencies in a time series affect the likelihood of a short time making a spurious conclusion about how a process varies in time. The tool then compiles typical statistics used to describe the relationships in the sampled data, through repeated samplings, and then use these derived data to gain insights to the questions, *how often are the trends observed in short-term data misleading, and can the characteristics of these trends be used to predict our likelihood of being misled?* Findings from this work will support the development of a deep understanding of temporal scaling in ecology, aiding in the interpretation of countless future short-term studies. Secondly, and more broadly, our findings have applicability across a variety of domains. Results from this approach will have the opportunity to guide science funding policy, experimental design and interpretation, and data archiving.

## 2 Materials and Methods

### 2.1 Developing the ‘broken_window’ analysis algorithm

The broken_window algorithm breaks a time series dataset into all possible sequential subsets and then fits a linear model to each of these subsets and compiles the resulting summary statistics, allowing a user to identify and quantify spurious trends within their data. The algorithm is implemented as a series of functions written in R. The algorithm requires a user-inputted two variable data frame with a regular measurement interval as the first variable, and a response variable as the second variable. For the purpose of this study, we assume a yearly measurement interval and some integrative response metric (captures of organisms per trap, average reading, total yield). Data are first subjected to a standardization function which converts the response metric to a unitless Z-score to normalize the data and make it possible to compare datasets with responses of very different magnitudes, and to minimize the impact of measurement unit choice on the observed trends.

A function that fits a linear model to the data and computes an output vector with the number of observations, the number of years in the study, and particular summary statistics of interest, namely, the slope of the relationship between the response variable and time, the standard error of this relationship, p-values for each of these statistics, and then *R*^*2*^ and adjusted *R*^*2*^. Although *R*^*2*^ and *p* are not measures of statistical confidence per se, they are often used by ecologists in this way (Nakagawa & Cuthill, 2007; Yoccoz, 1991), and thus can be used as a means to approximate ‘conclusions’ that a researcher might make of the data. We use this fitting function within a moving window function that takes a provided data frame and iterates through it at all possible subsets and intervals, feeding each interval to the fitting function described above, and compiling the fit statistics for each into a single object.

## 3 Calculation

The moving window function is defined as follows. Let ***D*** represent the complete dataset, with ***D***_***t***,***r***_ representing a single observations of time ***t*** and response ***r***. Let ***Y = (y***_***1***_, ***y***_***2***_, ***…, y***_***n***_) represent the set of unique values of ***t*** for which observations are recorded, where ***n*** is the total number of unique values of ***t. D*** is partitioned into sequential subsets of size ***S = (3, 4, …, n)*** to create windows ***w***_***Y***,***S***_ such that each window

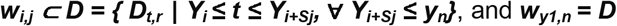

For each ***w***_***i***,***j***_, we apply the fitting function described above, and compile the resultant fit statistics for downstream analyses into a data frame. Then, we calculate several meta-statistics and produce visualizations of trends from the resultant data frame.

First, we defined the slope of the longest time series (i.e. the slope of the linear regression of the whole dataset, ***D***) as the proxy for the ‘true’ trajectory of the data (as it represents the best information available), along with the computed slope’s standard deviation and standard error of the mean as measures of the ‘true’ variability of the set. Meta-statistics are computed based on comparison to these ‘true’ statistics.

For all meta-statistics based on frequentist assumptions, we used a set of frequently used ‘significance’ levels as defaults (i.e. an α=0.05 for line fit statistics) but also encoded the functions so that a user could change these default values easily through supplying a function with different arguments. For each relevant function, we allowed users to toggle via a function argument between these meta-statistics based on the full set of windows tested, or only on the set of windows with statistically significant results, as defined above.

We defined “stability time” as the number of time steps needed before a given proportion of slopes (default = 95%) observed in a window of that length are within a certain number of standard deviations (default = 1) of the true slope. These values were selected to mitigate the impact of outlying data and to reflect industry standards. We computed absolute range (minimum and maximum values) of slope across all windows, as well as relative range (minimum and maximum difference from the ‘true’ slope, computed as the slope(***w***_***i***,***j***_) minus slope(***D***)). We also created functions that computed the proportion of windows examining a dataset would produce particular results. The proportion of statistically significant slopes produced by a given ***D*** measure the probability that a randomly selected window of time would produce a ‘statistically significant’ result. We defined the ‘proportion wrong’ as the proportion of windows producing statistics that would lead to conclusions differing from those observed for the ‘true’ trend (i.e. if the true trend was a positive slope, all windows suggesting a negative or non-significant zero-magnitude slope were considered spurious, and so on). We provide functions to compute the proportion wrong for all windows in combination, for each window length, and in the set of windows with lengths less than stability time. In combination, these functions provide a standardized approach to asking the questions of how long a system must be observed to make consistent conclusions about its trajectory, and the likelihood of coming to misleading conclusions about a system if it is observed for less than that time period.

We created several visualization functions to enable a user to, for a given dataset ***D***, quickly interpret trends based on these meta-statistics, and compare trends in outputs across multiple datasets. A pyramid plot (**Fig. 2A**) uses the data frame of summary statistics from the fits of all windows. It plots the computed slope for each window on the x axis and the length of the window on the y-axis, resulting in a triangular or funnel shaped cloud of points. By default, point size is scaled by the R^2^ of the response-by-time relationship within a given window and statistically significant points are demarcated by a circle, and non-significant points given by an ‘X’. All points are given with lines indicating their respective standard error. A vertical dashed line indicates the slope of the longest time series, and two dotted vertical lines are plotted at one standard deviation from this value, allowing a user to visually identify the stability time, that is, the length of time required for the majority of windows to produce slopes within a certain interval of the true slope.

**Figure 2:**
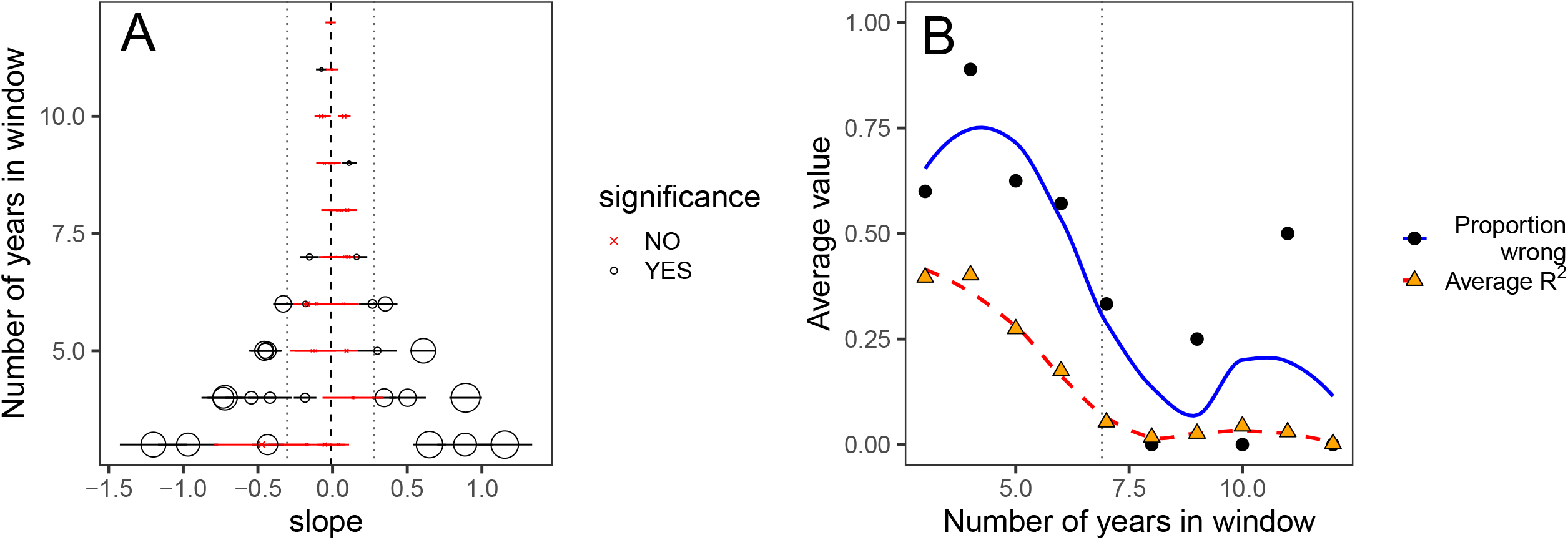
Core outputs of the broken_window algorithm: Using the firefly data from the early sucessional plant community presented in Hermann et al (2016), we are able to compile 55 possible windows of three years or greater. **A)** The pyramid plot gives a distribution of possible conclusions. On this plot, each point represents a window and its corresponding summary statistics for a linear relationship between the response variable (in this case, z-scaled population density of fireflies) and time. Point coordinates are defined by the slope and length of a window, and point size is scaled by the R^2^ computed for that regression. The lines accpmanying each point represent standard error of the slope for each point. Statistically significant relationships (in this case α=0.05) are plotted as black circles, and non-significant slopes are plotted as red Xs. The vertical central dashed black line represents the slope of the complete time series (here with 12 years of data) and the vertical dotted grey lines are placed at one standard deviation in both the positive and negative direction from the ‘true’ slope. **B)** The ‘wrongness plot’ visualizes the relationship between the likelihood of a spurious conclusion and statistical proxies for ‘confidence’ in a relationship. The proportion of windows where spurious slopes were observed by the length of window are displayed as black circular points with blue solid smoothing line, and the average R^2^ value across windows of that length are given as orange triangular points with a dashed red smoothing line. The grey dotted vertical line is placed at the ‘stability time’ of 7 years, after which the slopes in 95% of the windows occur within one standard deviation of the ‘true’ slope.

The “wrongness” plot (**Fig. 2B**) examines the same data from a summarized perspective-it plots the average R^2^ value and proportion wrong on the y axis by number of years in a window on the x-axis, allowing a user to visualize the relationship between misleading results and the ‘confidence’ in them for a given ***D***. Finally, the “broken stick” plot (**Fig. 3**) allows a user to visualize the raw time series from ***D*** simultaneously with some of the results of the broken_window algorithm. The z-scaled response metric (y-axis) is plotted by observation time (x-axis). The true slope of the entire dataset ***D*** is plotted as a solid black line. Then, best fit lines for each window of a user-specified length (default=3-time steps) are plotted, allowing a user to visualize the variation in trend at different points in the time series. Statistically significant slopes are given by dashed red lines, non-significant slopes are indicated by dotted lines. Finally, we created a function which layers and animates broken stick plots to visualize how window slopes change given increasing window length.

**Figure 3:**
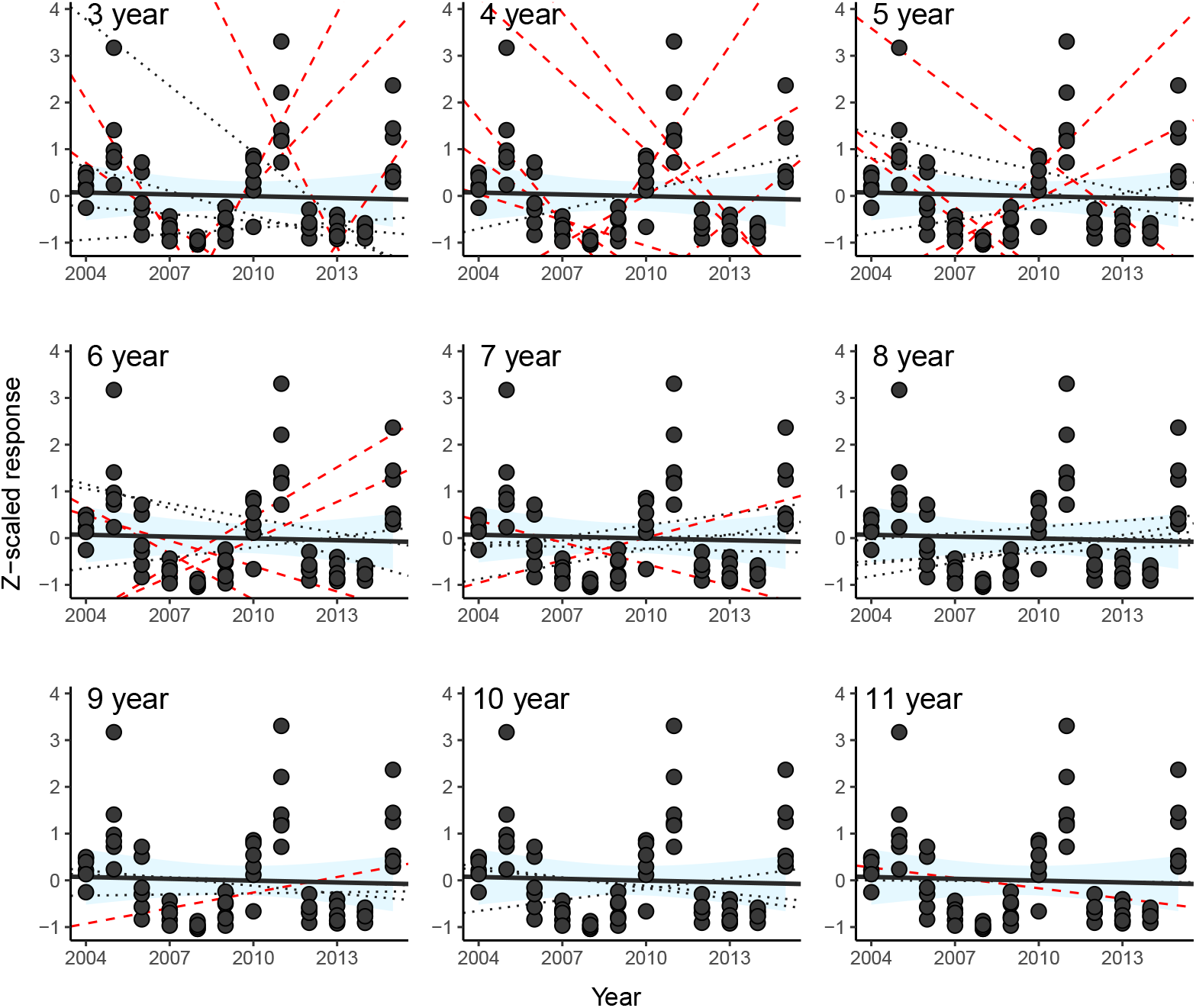
The broken stick plot allows a user to visualize the magnitude of difference between the slopes produced at different window lengths. Using the firefly data from the early successional plant community from Hermann et al (2016), all of the nine panels presents the Z-scaled response of firefly density over time, and a solid black line indicates the linear regression of the full data series (the ‘true’ slope). The 95% confidence interval of this line is plotted in light blue. Within each panel, the linear regressions for each window of a given length are plotted: regressions with a statistically significant slope (at α=0.05) are given with red dashed lines, and non-significant regressions are plotted as grey dotted lines.

The R script was developed in RStudio Version 1.2.5033 “Orange Blossom” running R 3.6.2 “Dark and Stormy Night.” The script, its development history and all code for case studies and figure generation, are available on GitHub at https://github.com/cbahlai/broken_window.

## 4 Results

We demonstrated the utility of the broken_window algorithm using the firefly study which inspired its development (Hermann et al., 2016). These data on firefly (beetles in the family Lampyridae, with those captured primarily thought to belong to *Photinus pyralis*) captures on insect sticky traps were collected 2004-2015 across 10 plant communities in southwestern Michigan. Complete sampling design and treatments descriptions are provided in Hermann et al (2016). For the purpose of this demonstration, we used the data collected at the perennial early secessional community plots, where fireflies were relatively abundant and complete data were available. Data were subjected to cleaning and quality control using scripts developed by Hermann et al (2016), and then compiled into a metric of total captures per trap, by year (N=12) and replicate (N=6), for a total of 72 observations.

The broken_window algorithm produced 55 unique windows (1 sequence of 12 years of data, 2 sequences of 11 years of data, …, 10 sequences of 3 years of data). The full 12 year, 72 observation dataset of the normalized response over time was found to have a non-significant slope (−0.01±0.03, p=0.70) and low R^2^ value (0.002) suggesting there is unlikely to be a linear trend with time in these data (or, more specifically, we fail to reject the null hypothesis that there is no linear relationship between our response and time) (**Fig. 2A**). Values computed for the slopes across the various windows ranged ±1.2 units around the true slope. The algorithm found a stability time of 7 years, that is, once seven years of data were collected, slopes on >95% of windows tested from anytime in the study were within one standard deviation of the slope of the longest series. Overall, nearly half (27/55) of the windows tested found a statistically significant slope, and thus there was nearly a 50% chance a shorter sample leading to a misleading conclusion. Although misleading slopes combined with significant p-values occurred for window lengths longer than 7 years (**Fig. 2B**), they were much more common with window lengths shorter than the stability time (68% of windows), yet these shorter windows were also more likely to be accompanied by a R^2^> 0.1 (**Fig. 2B**). Although 3 of these 21 windows ≥7 years in length contained statistically significant trends, after stability time, relative slope ranged from -0.14 to 0.17 z-scaled units around the true slope (**Fig. 3**).

## 5 Discussion

Patterns observed in local scale, short-term ecology tend to be dominated by stochastic forces, making generalizations, extrapolations and predictions difficult at larger scales, yet are essential to capture fine-scale understanding of system dynamics (Chave, 2013; Willis & Birks, 2006). The broken_window algorithm formalizes a framework for determining how long a system must be observed before conclusions about its general trends can be reached, and the prevalence of misleading results that occur prior to that time period. With our firefly case study, we found that trends observed prior to our ‘stability time’ of seven years had essentially even odds of being misleading: of three possible outcomes for each window (slope more negative than overall trend, slope more positive than overall trend, slope the same as overall trend), 2/3 of outcomes fell into the two former, and erroneous categories. In this case, no net linear trend was observed in the firefly population data (**Fig. 1, 2A**). Interestingly, we observed that in our case study, statistics commonly used as indicators of “strength” of relationship suggested more uncertainty, and less ‘confidence’ in results from windows of longer length: p-values, on average, went up, and R^2^ values decreased on average as longer windows of the time series were examined (**Fig. 2C**). This finding shines an important light on the reliability of these statistical tools as indicators of model performance: although they provide measures of how well the data from a given window fit the selected model at that time, they also inflate our confidence in what is often an inappropriate model fit to a spurious or short-term trend (Nakagawa & Cuthill, 2007). Given the high likelihood that these observations will vary by context, future work must consider how process characteristics, data availability, and cultural precedent (i.e.: the history of use of a given approach in a scientific field) affect the selection and interpretation of these models. Furthermore, it should explicitly examine data with different structures to examine the relationship between time series shape and likelihood of erroneous conclusions at differing study lengths.

In this paper, we demonstrate the utility of the broken_window algorithm in the context of a simple, single population case study. However, this analytical approach has broad application which has been applied by several colleagues in additional systems. In a recent study, Cusser et al (2020) applied the algorithm to a thirty-year experiment comparing the sustainability and productivity attributes of an agricultural cropping system under several management regimes. In this system, due to high variability between treatments, 15 year observation periods were needed to detect consistent between-treatment differences in yield and soil water availability, and at least 1/5 of all windows examined resulted in spurious, statistically misleading trends (i.e. suggest the opposite relationship between management treatments). In an expansion of this work, Cusser and colleagues (2021) used the broken window algorithm to mine more than 100 additional long-term population datasets and found that ∼50% of studies had temporal dependencies between treatments that could not be reliably detected with fewer than 10 years of data. Furthermore, they linked the stability of the abiotic environment to the ability to detect trends: simply, experiments taking place in more variable environments were more prone to spurious trends and required more data and time to establish experimental differences. In another study, R. Christie et al (2021) compiled 289 surveys of deer tick activity produced by public health departments and researchers primarily in the northeast and Midwest United States and subjected each set of observations to the broken_window algorithm. They found none of the survey data reached stability time in less than 5 years, indicating that shorter term studies may be insufficient to infer long term population dynamics. Bruel and White (2021) used a similar approach to investigate the optimal sampling of sediment cores for constructing phytoplankton communities. However, they examined the sampling effort required to detect abrupt shifts (i.e., changepoints) in community structure, as opposed to simple linear trends over time. This work highlights the need for future studies investigating the sampling required to detect patterns beyond those from simple linear regression. Other related work has focused on estimating the length of time series required to achieve high statistical power (White, 2019), and studying data-poor fisheries (White & Bahlai, 2020): taken together, these tools will enable previous work to be mined to understand the characteristics of trends common to those systems, and enable future studies to be designed to maximize information value.

The broken_window algorithm uses the longest available study duration as a proxy for ‘truth’ as its core assumption. However, long-term studies themselves are not immune to uncovering misleading trends. Methodology, site selection, and periods of disturbance following the initiation of a long-term study may inherently bias the apparent trajectory of a system (Fournier et al., 2019). This highlights the importance not just of study duration, but of the selection of study starting and ending points: capturing an outlying data point or a high or low in a system’s natural variability near the beginning or end of the study period will be highly influential on the statistical outcome, and thus the conclusions reached (Chatterjee & Hadi, 1986; Fournier et al., 2019). Understanding and characterizing these highly influential observations in the analysis process is essential to our interpretations of these ecological trajectories. Thus, it is important to consider these biasing factors when using long-term data in algorithms like the one presented herein: any statistical method is likely to be influenced by outlying or unlikely observations.

The broken_window algorithm uses a linear model as its underlying structure, which is the simplest case of a relationship a response variable might take with time. However, many ecological processes are not linear with time and may be better described with non-linear approaches (Bahlai & Zipkin, 2020; Knape, 2016; Wauchope et al., 2019). In the initial deployment of this algorithm, we created a tool for the simplest case that would be applicable under a whide variety of circumstances, but future iterations should consider multiple underlying model structures, as well as contingencies for unevenly spaced observations or missing data.

## 6 Conclusions

The ever-increasing availability of long-term data, fostered by the growth of technology that enables automated collection and sharing of data products, and the infrastructure availability and ‘maturity’ of projects like the US (and international) Long Term Ecological Research networks (Brunt et al., 2002) and more recently, the National Ecological Observatory Network (SanClements et al., 2020; Schimel et al., 2007) present several key opportunities for new understanding of temporal processes in ecology. Not only can these data be used to observe long-term processes in their respective systems, these data can be used to contextualize the vast amount of data produced by shorter-term studies in our field. Ecology, until relatively recently, was a field defined by data scarcity: studies took place at local scales, over time periods manageable to small groups of researchers, and these shorter-term studies remain the most common output in ecological research (Peters, 2010). Their work represents a huge human undertaking, however, and it is critical that we are able to interpret the insights these observations provide appropriately.

The broken_window algorithm provides a framework for understanding how ecological data produced by different domains behaves at different temporal scales. Thus, this tool can be used to synthesize data describing ecological processes, specifically examining how system properties (such as landscape, site, seasonality, lifespan in the case of organisms, management regimes, cycles in population trends) affect the likelihood of a spurious trend being observed. In future work, we will examine data of differing structures to identify the characteristics of observation periods that are more likely to produce misleading results, and conversely, the characteristics of time periods that are consistent with longer system trends. This framework will support ongoing research efforts to separate trends in ecological systems from natural variability, human biases and research-specific influences and underlying processes, and provide critical insight into the scaling to temporal processes between short- and long-term experimental designs.

## Acknowledgements

Data used in our firefly case study was collected on traditional Anishinaabe land where Hickory Corners, Michigan is currently located. The broken_window algorithm was initially inspired by conversations with John Andrew Gerrath, Ilya Gelfand, and Doug Landis and through feedback and refinements from G. Phillip Robertson, Scott Swinton, Elise Zipkin, Nick Haddad, and the rest of our colleagues at Kellogg Biological Station. Additionally, the algorithm has incorporated feedback from colleagues from the United States Long Term Ecological Research network throughout its development. A particular thanks to Sven Bohm for database curation. Infrastructure supporting this work was funded by the National Science Foundation Long-term Ecological Research Program (DEB 1832042) at the Kellogg Biological Station; the algorithm was developed with funding from the National Science Foundation Directorate for Computer and Information Science and Engineering (OAC 1838807) to CB, JP and KSW and was completed with the support of a grant from the National Science Foundation Directorate for Biological Infrastructure (IIBR 2045721) to CB.

